# Oncogenic KRAS Requires Complete Loss of BAP1 Function for Development of Murine Intrahepatic Cholangiocarcinoma

**DOI:** 10.1101/2021.10.12.464103

**Authors:** Rebecca Marcus, Sammy Ferri-Borgogno, Abdel Hosein, Wai Chin Foo, Bidyut Ghosh, Jun Zhao, Kimal Rajapakshe, James Brugarolas, Anirban Maitra, Sonal Gupta

## Abstract

Intrahepatic cholangiocarcinoma (ICC) is a primary biliary malignancy that harbors a dismal prognosis. Oncogenic mutations of *KRAS* and loss of function mutations of BRCA1-associated protein 1 (*BAP1*) have been identified as recurrent somatic alterations in ICC. However, an autochthonous genetically engineered mouse model of ICC that genocopies the co-occurrence of these mutations has never been developed. By crossing *Albumin*-Cre mice bearing conditional alleles of mutant Kras and/or floxed *Bap1*, Cre-mediated recombination within the liver was induced. Mice with hepatic expression of mutant *Kras*^G12D^ alone (KA), bi-allelic loss of hepatic *Bap1* (B^homo^A), and heterozygous loss of *Bap1* in conjunction with mutant *Kras*^G12D^ expression (B^het^KA) developed primary hepatocellular carcinoma (HCC), but no discernible ICC. In contrast, mice with homozygous loss of *Bap1* in conjunction with mutant *Kras*^G12D^ expression (B^homo^KA) developed discrete foci of HCC and ICC. Further, the median survival of B^homo^KA mice was significantly shorter at 24 weeks, when compared to median survival of ≥40 weeks in B^het^KA mice and approximately 50 weeks in B^homo^A and KA mice (p <0.001). Microarray analysis performed on liver tissue from KA and B^homo^KA mice identified differentially expressed genes in the setting of BAP1 loss and suggests that deregulation of ferroptosis might be one mechanism by which loss of BAP1 cooperates with oncogenic Ras in hepato-biliary carcinogenesis. Our autochthonous model provides an *in vivo* platform to further study this lethal class of neoplasm.

## Introduction

Cholangiocarcinomas are a group of primary liver cancers arising from the biliary epithelium. These aggressive cancers are the second most common primary hepatic malignancy after hepatocellular carcinoma (HCC), and they currently represent 15% of all primary liver tumors and 3% of gastrointestinal cancers [1]. Traditionally, cholangiocarcinomas are classified based on anatomic site of origin and include carcinomas arising from the intrahepatic, perihilar, and extrahepatic biliary tree [2]. While the incidence of perihilar and extrahepatic disease has been decreasing [3–5], the incidence and mortality related to intrahepatic cholangiocarcinoma (ICC) has been rising over the past several decades in many parts of the world [1,5]. This is perhaps most notable within the Western Hemisphere [6–7], as there has been a doubling of the annual incidence of reported ICC in the United States since the 1970s [8]. This trend was noted to have accelerated during the last decade, with an annual percent increase of >4% [8].

At present, the only potential curative treatment for ICC is surgical resection; however, the majority of patients have advanced disease at the time of diagnosis [9], and ultimately only 15% of patients are eligible for curative resection [10–11]. Furthermore, amongst patients undergoing curative-intent resection, treatment failure is common, with recurrence in as many as 66% of patients and median overall survival of only 28-36 months [10,12]. Unfortunately, despite the advances made in available systemic and targeted therapies for many malignancies, the available treatment options for patients with ICC remain limited [13–16]. Moreover, best practice recommendations continue to be debated [17] with respect to which patients should receive systemic therapy, the therapeutic regimen they should receive, timing of systemic therapy in relation to surgical intervention, and the use of associated radiotherapy [1,9,18–20]. This translates into the poor prognosis associated with ICC, with <5% of patients being alive 5 years after diagnosis [10,21–22].

Despite its dismal prognosis and rising incidence, there are still substantial deficits in our current understanding of ICC, including the mechanisms of biliary carcinogenesis. This is at least in part due to an incomplete assortment of experimental tools for studying ICC, including a paucity of available cell lines and faithful genetically engineered mouse models (GEMMs). Recent molecular studies of ICC tumors have identified recurrent somatic mutations in several oncogenes and tumor suppressor genes. For example, oncogenic *KRAS* mutations have been observed in 5-27% of ICCs [23–24]. This is not unexpected given that Ras signaling has been demonstrated to be deregulated in numerous human tumors, and that *KRAS* is the most commonly mutated oncogene in human cancers [24–25]. Loss-of-function mutations of the *breast cancer type 1 susceptibility protein (BRCAl)-associatedprotein 1* (*BAP1*) have also been identified in 7-32% of Western ICCs [23]. *Bap1* mutations are also seen in Eastern ICCs, which are frequently associated with liver-fluke infections, albeit at slightly lower frequencies than seen amongst Western ICCs [24]. Previous studies have suggested that BAP1 protein functions as a tumor suppressor [27–28] and both germline and somatic mutations of *BAP1* have been identified in numerous tumor types, including melanoma, mesothelioma, renal cell carcinoma, and breast cancer [29–32]. BAP1 is a nuclear deubiquitinating enzyme in the ubiquitin carboxyterminal-hydrolase subfamily that is involved in chromatin remodeling [27]. Loss of BAP1 links deregulation of the cell death mechanism known as ferroptosis to carcinogenesis. Indeed, recent studies have demonstrated that BAP1 represses expression of the cystine transporter SLC7A11 and, as a consequence, inhibits cystine uptake, leading to elevated lipid peroxidation and ferroptosis-mediated cell death. Cancer-associated *BAP1* mutants lose their abilities to repress SLC7A11, leading to attenuation of ferroptosis and tumor promotion [33–34].

In this study, we sought to investigate the cooperation between Ras and BAP1 in ICC pathogenesis by generating autochthonous mice with *Cre*-mediated conditional activation of mutant *Kras* and/or deletion of *Bap1* alleles within the Albumin (*Alb*) expressing domain, which is comprised of liver progenitor cells, as well as adult hepatocytes and cholangiocytes [35–36]. Cohorts of mice with oncogenic *Kras* activation alone (KA), bi-allelic loss of hepatic *Bap1* (B^homo^A), or the expression of oncogenic *Kras* activation in combination with heterozygous *Bap1* loss (B^het^KA) developed HCC with variable penetrance after a prolonged interval, but none of the mice developed ICC. In contrast, the expression of mutant *Kras* in conjunction with bi-allelic *Bap1* deletion (B^homo^KA) resulted in development of ICC that recapitulated the histological features of human ICC, albeit admixed with foci of HCC in the liver parenchyma. Thus, our findings demonstrate that complete abrogation of BAP1 function might be one of the requirements for developing an ICC phenotype in mice expressing oncogenic Kras in the Alb-expressing domain, underscoring the importance of this tumor suppressor gene in ICC pathogenesis. Further, transcriptomic profiling and immunohistochemical validation confirmed upregulation of the cystine transporter xCT (encoded by *SLC7A11*) in B^homo^KA compared to KA neoplasms, suggesting a potential role for ferroptosis deregulation in the tumor promoting phenotype induced by BAP1 loss. This work establishes a relevant autochthonous model of ICC that genocopies the co-occurrence of two recurrent mutations observed in a subset of human ICC, and provides an opportunity to evaluate putative actionable pathways against this lethal disease in an *in vivo* setting.

## Materials and Methods

### Generation of conditional BAP1 mice

BAP1^L/L^ mice were kindly provided by Dr. James Brugarolas (University of Texas Southwestern Medical Center) and are now available commercially (Stock No: 031565. Jackson Laboratories, Bar Harbor, Maine, USA) [37–38]. *Albumin*-Cre mice have been previously described [35–36] and were purchased from Jackson laboratories (Stock No: 016832). Lox-STOP-Lox-*Kras^G12D^* mice have also been described before [40] and were also purchased from Jackson laboratories (Stock No: 019104). BAP1^L/L^ mice were crossed with Alb-Cre or Albumin-Cre;LSL-Kras^G12D^ mice to generate Albumin-Cre; BAP1^L/+^ (B^het^A), Albumin-Cre; BAP1^L/L.^(B^homo^A), Albumin-Cre;Kras^G12D^; BAP1^L/+^ (B^het^KA), and Albumin-Cre,Kras^G12D^; BAP1^L/L^ (B^homo^KA) mice. All mice were housed in a pathogen–free barrier facility with food and water *ad libitum*. Animal studies were conducted in compliance with Institutional Animal Care and Use Committee (IACUC) guidelines of the University of Texas MD Anderson Cancer Center and performed in accordance with the NIH guidelines (https://grants.nih.gov/grants/olaw/guide-for-the-care-and-use-of-laboratory-animals.pdf) for use and care of live animals under the protocol number 00001937-RN00 (expiring May 2022). PCR was performed to confirm the genotype of mice using DNA obtained from tails.

### Genotyping PCR

DNA extraction from mouse tail clips obtained at 7-10 days of age was performed using REDExtract-N-Amp™ Tissue PCR Kit (Cat#R4775, Sigma Aldrich) following the manufacturer’s protocol. In brief, tail clips were combined with 100 μL of Extraction Solution and 25 μL of Tissue Preparation Solution and mixed by vortexing. Samples were incubated for 15 minutes at room temperature followed by incubation at 95°C for 5 minutes. 100 μL of Neutralization Solution was added to each sample and again mixed by vortexing. Extracted DNA was subsequently used to perform genotyping PCR. Primer sequences and expected amplicon sizes are found in Supplementary Figure 1.

### Histology and immunohistochemistry

Mouse tissue obtained during necropsies were fixed in 10% neutral buffered formalin for 72 hours. They were subsequently processed, embedded in paraffin, sectioned, and stained with hematoxylin and eosin (H&E) and reticulin. For immunohistochemistry, 5 μm sections were incubated with primary antibodies as described previously [41]. For analysis of marker expression at least three mice per genotype were characterized, and ImageJ software was used for quantification of tissue sections. Antibodies used are as follows: Anti-BAP1 (Cat#PA5-12061, ThermoFisher Scientific), anti-CK-19 (Cat#MABT913, Millipore Sigma), anti-HepPar1 (Cat#NBP2-45272, Novus Biologicals), and anti-SLC7A11 (Cat#NB300-218, Novus Biologicals). ImmPRESS-HRP-linked anti-rat (Cat#MP744415) was purchased from ThermoFisher Scientific and Mouse on Mouse ImmPRESS-HRP-linked anti-mouse (Cat#MPX-2402-15) from Maravai LifeSciences.

### Microarray analysis

After histopathologic confirmation of the presence of primary liver tumors, total RNA was extracted from three KA and three B^homo^KA flash-frozen tumors using the RNeasy Mini kit (Cat#74106, Qiagen) according to the manufacturer’s instructions. Samples were submitted to the Advanced Technology Genomics Core (ATGC) facility at the University of Texas MD Anderson Cancer Center. Briefly, the concentration of total RNA was assessed using the Nanodrop ND-1000 Spectrophotometer (ThermoFisher Scientific). Once the sample concentration was determined, the integrity of the total RNA was assessed using the Agilent 2100 Bioanalyzer Nanochip assay. Samples with a minimum concentration of 33 ng/ul were selected for target amplification with the Whole Transcript (WT) Plus assay (Affymetrix). 100 ng of total RNA was used to process the samples for Whole Transcriptome Expression Analysis with the WT Plus assay.

Samples were reversed transcribed to generate amplified, fragmented, and biotinylated sense-strand cDNA (sscDNA), according to manufacturer’s standard protocol. 5.2 μg of fragmented and labeled sscDNA was then hybridized to Affymetrix Mouse Transcriptome Array 1.0 ST at 45°C for 16 hours and subsequently washed and stained using Affymetrix proprietary reagents in the GeneChip Fluidics Station 450 and scanned in the GeneChip Scanner 3000 7G (Affymetrix).

CEL files generated after Mouse Transcriptome 1.0 ST GeneChip scanning were uploaded onto the Expression Console software. The CEL files were pre-processed and normalized using the Robust Multichip Average (RMA) algorithm implemented in oligo [42] package within R statistical software (R, v4.0.2). Differentially expressed genes between the experimental groups were identified using the Bioconductor limma [43] package imposing a filtering criteria of fold change >2 (<0.5) and p <0.05. Clustering and heatmaps were generated using Matplotlib, NumPy and SciPy libraries under python. Gene Set Enrichment Analysis (GSEA) using Molecular Signature Database (MSigDB) [44] was performed to find enriched pathways (q <0.25).

### Statistical analysis

Statistical analyses were performed using Prism 8 (GraphPad, San Diego, CA, USA). Statistical significance was determined using the unpaired Student’s *t* test with Welch’s correction and two-way ANOVA with Sidak’s post hoc test, as appropriate. For all experiments with error bars, standard deviation (SD) was calculated to indicate the variation within each experiment and data, and values represent mean ± SD. Kaplan–Meier method and log-rank test was used for survival analysis of mice. A p-value of <0.05 was regarded as statistically significant.

## Results

We generated a genetically engineered mouse model (GEMM) with a conditional knockout allele for *Bap1* (BAP1^L/L^) and a conditionally activated allele for *Kras* (LSL-Kras^G12D^) (Figure 1a). In our cohorts of GEMMs, Cre expression was under the control of the *Albumin* promoter, resulting in expression of oncogenic *Kras* and loss of *Bap1* in liver progenitor cells during late embryogenesis, as well as within Alb-expressing hepatocytes and cholangiocytes of adult mice [36]. Of note, several previously described ICC GEMMs have utilized the *Albumin* promoter to target genetic alterations of interest to the liver [35,45–47]. The *Bap1*^L/L^ allele sustains Cre-mediated excision of exons 4 and 5, resulting in a functionally inactive gene as previously described [48] (Figure 1a). The *LSL-Kras^G12D^* allele results in oncogenic *Kras* expression at endogenous levels following Cre-mediated excision of a transcriptional stop element [25]. Compound mutant mice with the following genotypes were achieved through multiple generations of crossbreeding: Alb-Cre;Kras^G12D^ (KA), Alb-Cre;BAP1^L/L^ (B^homo^A), Alb-Cre;Kras^G12D^;BAP1^L/+^ (B^het^KA), and Alb-Cre;Kras^G12D^;BAP^L/L^ (B^homo^KA) (Figure 1a).

**Figure 1:**
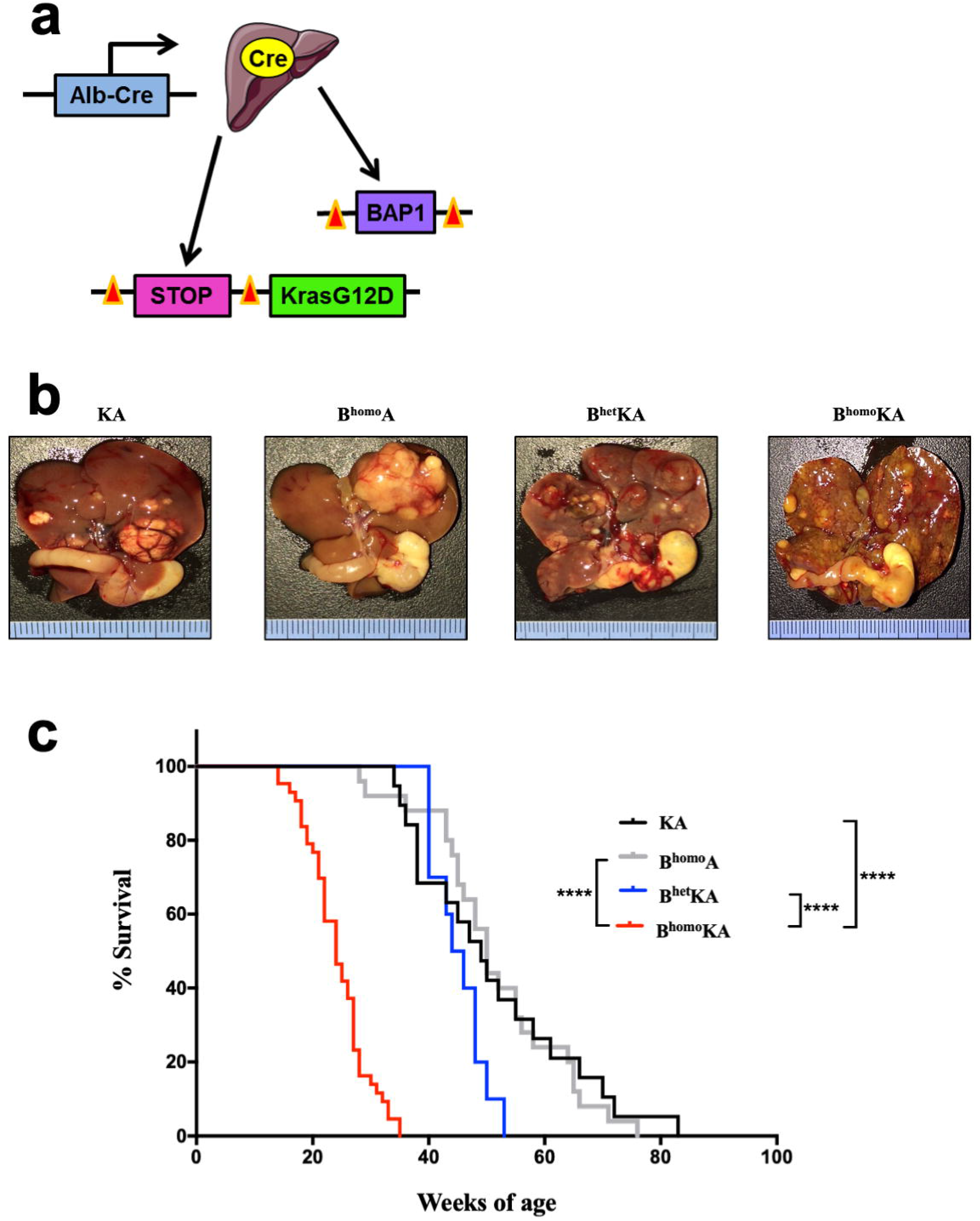
Mice with constitutional KrasG12D activation and BAP1 deletion in the hepatic epithelium develop primary liver tumors. (a) Modeling strategy to generate compound mutant mice. Mice harboring *Albumin*-Cre transgene, Lox-STOP-Lox-*Kras*G12D, and BAP1^L/L^ were created to conditionally activate KrasG12D and delete BAP1 in the hepatic epithelium; (b) In situ gross tumor nodules throughout the hepatic parenchyma with associated hepatomegaly in experimental cohorts; (c) Kaplan–Meier survival analysis for KA (N=17), B^homo^A (N=25), B^het^KA (N=10), and B^homo^KA (N=43) cohorts. Two-tailed unpaired Student’s t test was used for data analysis and considered significant if ****, p < 0.0001.

### *Tissue specific concomitant expression of oncogenic* Kras *and* Bap1 *loss results in primary liver tumors and decreased survival*

We first evaluated the gross intraabdominal findings and survival of all experimental cohorts. KA, B^homo^A, B^het^KA, and B^homo^KA mice were produced in the expected Mendelian frequencies. No cohort demonstrated evidence of early developmental abnormalities. Animals were monitored via serial examinations until they developed signs of illness including abdominal distension and bloating, cachexia, jaundice, diminished activity, and anorexia. Mice demonstrating one or more of these moribund criteria were euthanized, with the exception of those mice included for timed necropsies.

At the time of necropsy, all moribund animals were found to have hepatomegaly and solid liver tumors of various sizes located throughout the liver parenchyma (Figure 1b). These macroscopic findings corresponded to the abdominal distension observed on serial animal exams. Hepatic tumors presented as isolated nodules or, more often, as multiple independent lesions. Rarely, fluid-filled cystic lesions were found in addition to solid tumors. Some uncommon features observed at necropsy (all in less than 10% of mice) were areas of cystic degeneration or local invasion into adjacent organs (e.g., stomach, duodenum, pancreas and spleen).

The majority of KA mice lived for several months before exhibiting any signs of disease, after which they survived several additional weeks before requiring euthanasia for severity of symptoms (Figure 1c). The median survival of this cohort was 49 weeks (Figure 1c). The natural history of disease in the B^homo^A mice was comparable, with no significant differences in disease penetrance, timing of symptom onset, or rate of disease progression as compared to KA mice, with a median survival of 50 weeks (Figure 1c). The heterozygous loss of *Bap1* allele on a backdrop of oncogenic *Kras* expression in the B^het^KA mice resulted in acceleration of disease onset, with a median survival of 45 weeks, while the B^homo^KA mice demonstrated the most aggressive natural history amongst all genotypes, with a median survival of only 24 weeks (p <0.0001 for B^homo^KA versus other cohorts) (Figure 1c). Taken together, these data demonstrate that concomitant oncogenic *Kras* and homozygous deletion of *Bap1* results in reduced survival compared to the presence of either of these mutations individually or heterozygous deletion of *Bap1*.

### KrasG12D activation in combination with homozygous Bap1 deletion results in development of ICC

We next performed histopathologic analysis of the livers from each experimental cohort. Mice from all cohorts eventually developed hepatocellular carcinoma (HCC), albeit at different ages and with variable penetrance (Figure 2a, Table 1, Supplementary Figure 2). To confirm the effectiveness of *Bap1* gene deletion in B^homo^A, B^het^KA, and B^homo^KA mice, we evaluated BAP1 protein expression via immunohistochemistry (IHC) (Supplementary Figure 3). As expected, the B^homo^A and B^homo^KA mice show no BAP1 expression by IHC, while B^het^KA show decreased BAP1 expression as compared to KA control mice (Supplementary Figure 3). Of note, BAP1 expression is lost within both the normal hepatic parenchyma (Supplementary Figure 3a) and all primary liver lesions (i.e., ICC and HCC) (Supplementary Figure 3b).

**Figure 2:**
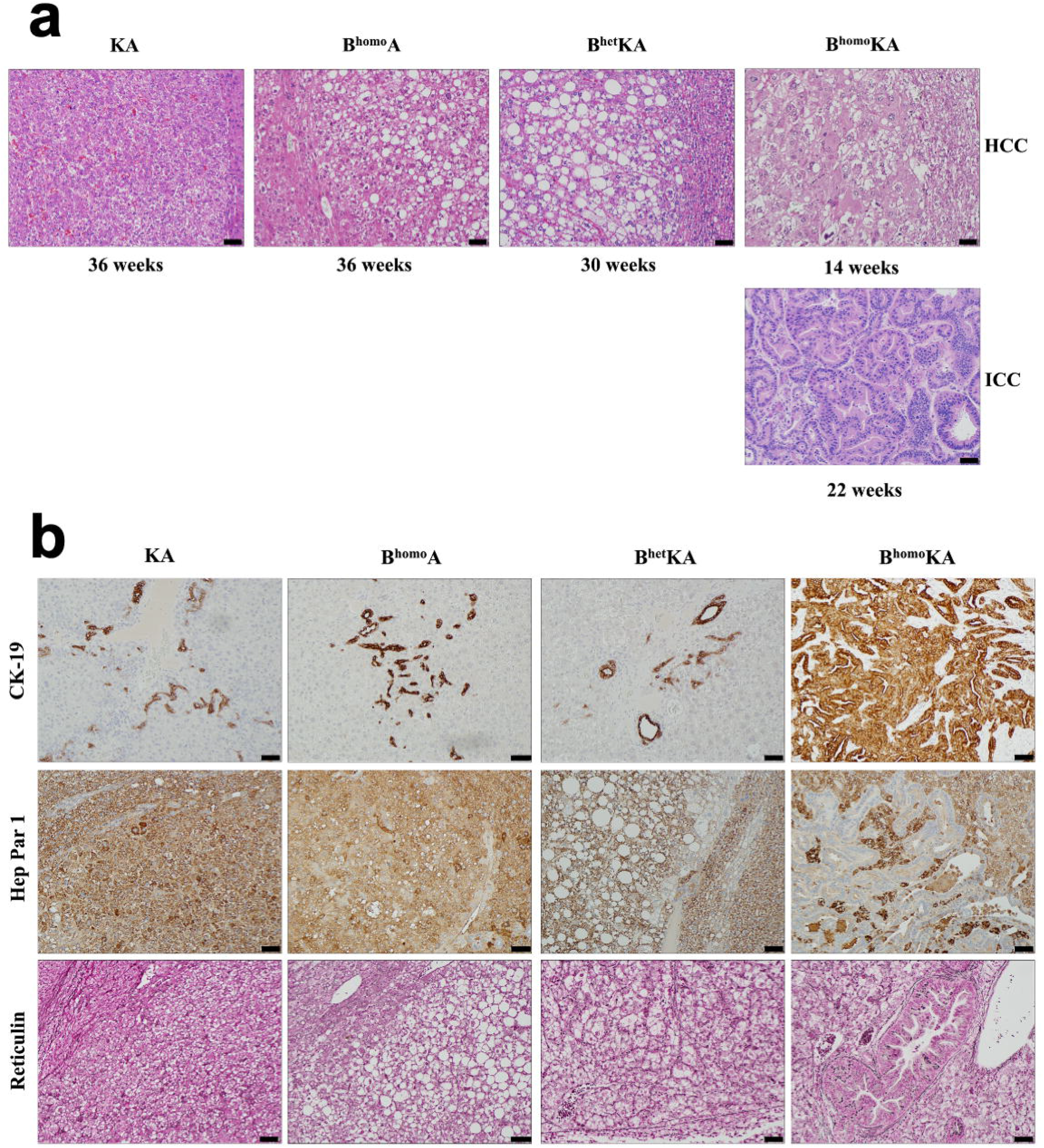
GEMM experimental cohorts have distinct histologic phenotypes. (a) KA, B^homo^A, and B^het^KA mice exhibit HCC only by H&E, while B^homo^KA mice have both ICC and HCC lesions; (**b**) CK-19, Hep Par 1, and reticulin staining demonstrates HCC lesions in all cohorts and both HCC and ICC in B^homo^KA mice. CK19 highlights normal ducts in KA, B^homo^A, and B^het^KA mice, while ICC foci are strongly positive in B^homo^KA mice. Conversely, Hep Par 1 stains positive amongst HCC foci in KA, B^homo^A, and B^het^KA mice, as well as in HCC foci of B^homo^KA mice; however, it is negative in ICC foci of the latter. Reticulin highlights abnormal architecture in HCC of all four mouse genotypes. Images were taken with 200x magnification. Scale bar is 50 μ*m*.

**Table 1.**
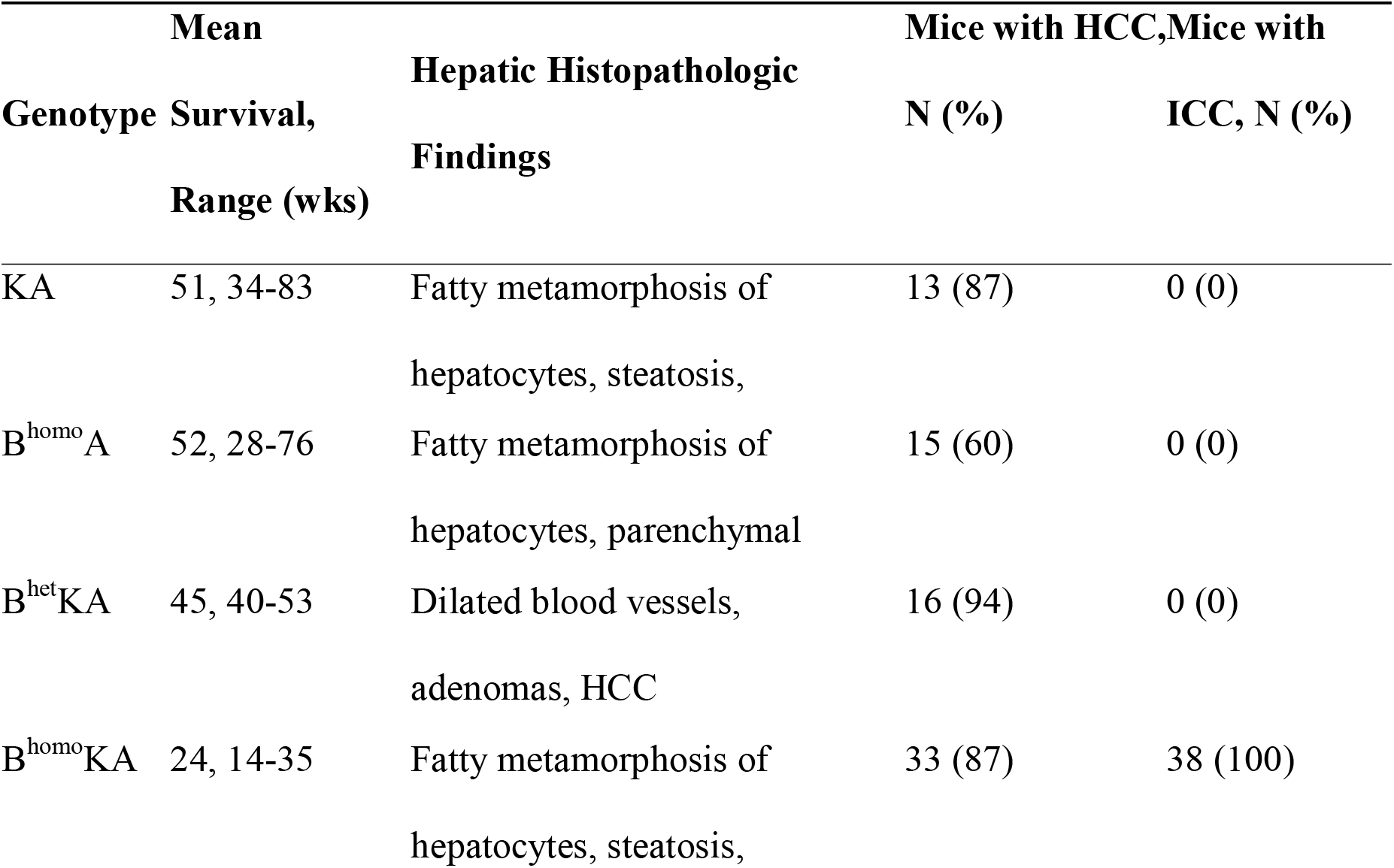
Hepatic genotypic and phenotypic differences between GEMM experimental cohorts. Total N includes necropsies performed as part of both survival analyses and timed necropsy analyses after the median survival of each cohort was reached.

In contrast to the other genotypes, microscopic analyses of the liver in B^homo^KA mice revealed a mixed phenotype consisting of both of ICC and HCC (Figure 2a). At 4-8 weeks of age, all experimental cohorts had normal liver histology (Supplementary Figure 2a-c) with the exception of B^homo^KA mice, in which bile duct hyperplasia was observed (Supplementary Figure 2d). Abnormal liver findings, including fatty transformation of hepatocytes, steatohepatitis, and parenchymal congestion, were eventually observed in all mice. Specifically, KA mice developed these changes between 20-28 weeks, B^homo^A mice between 24-32 weeks, B^het^KA mice between 20-24 weeks, and B^homo^KA mice developed these changes the earliest between 4-8 weeks of age (Supplementary Figure 2). The spectrum of hepatocellular disease included a variety of lesions, from hepatic adenomas to well-differentiated HCC to, in some cases, poorly differentiated HCC. The timing of development varied between experimental cohorts, with these pathologic findings occurring in B^homo^KA mice as early as 12 weeks of age and in KA mice as late as 40 weeks (Supplementary Figure 2). Furthermore, the proportion of mice within each cohort that developed HCC was lower in the B^homo^A mice (60%) mice than the other cohorts (87-94%, Table 1). The histologic spectrum of liver disease observed was similar across multiple generations of each experimental cohort.

Notably, histological findings of ICC were only observed in the B^homo^KA mice (Table 1). Specifically, microscopic changes consistent with bile duct dysplasia developed by 20 weeks of age, and foci of frank ICC were observed within 20-24 weeks (Supplementary Figure 2d). The degree of ICC differentiation varied between lesions, and even within the same lesion, but typically featured poorer differentiation in older mice. Overall, the timed necropsies show development of ICC only in mice with concomitant *Kras* activation and homozygous *Bap1* deletion. Furthermore, the earlier death exhibited in the B^homo^KA mice may be, at least in part, due to the development of the ICC.

To confirm the presence of two distinct differentiation lineages of neoplasm within our experimental cohorts, we stained tissue sections for markers of biliary epithelium (cytokeratin 19; CK-19) and hepatocytes (hepatocyte specific antigen; Hep Par 1) that have previously been used to distinguish ICC from HCC clinically [49–54]. Predictably, IHC analysis with CK-19 demonstrated robust staining of bile duct hyperplasia, dysplasia, and ICC in B^homo^KA mice, while only normal-appearing bile ducts stained positively with CK-19 in all other experimental cohorts (Figure 2b). Moreover, IHC for Hep Par 1 showed expression in hepatic adenomas and frank HCC in all experimental cohorts, while biliary epithelium-derived lesions did not show any staining (Figure 2b). Finally, staining with reticulin demonstrated a loss of the normal hepatic architecture in areas of hepatic adenomas and HCC development, but intact architecture within normal bile ducts, biliary hyperplasia and dysplasia, and frank ICC (Figure 2b). This is reminiscent of patterns seen in corresponding human disease processes where architectural disruption occurs with the development of hepatic adenomas and HCC, as well as other hepatocellular proliferative diseases, but not with ICC [55–57].

It should be noted that primary lung adenocarcinomas were observed in subsets of both KA and B^het^KA mice (Supplementary Figures 4). Specifically, 60% of KA and 55% of B^het^KA mice ≥24 weeks of age demonstrated lung lesions. These lesions were not seen in B^homo^KA mice, likely due to the accelerated natural history of the liver pathology in this mouse cohort. We confirmed by histology and CK-19 / Hep Par 1 IHC that the pulmonary lesions were primary lung adenocarcinomas, not metastatic HCC (Supplementary Figure 4). Notably, low levels of Albumin expression exists in the alveolar cells of adult lung per the Human Protein Atlas, and is also well reported in fetal lung tissues [58]. Therefore, the observed lung lesions likely represent breakthrough Cre expression in this organ.

### Transcriptomic profiling of B^homo^KA mouse tumors identifies potential effector pathways of tumor promotion caused by loss of BAP1

To further characterize the role of *Bap1* as a tumor suppressor gene, transcriptomic analysis was performed on harvested liver tumors from KA and B^homo^KA mice, with three mice per genotype used for microarray analysis. This analysis identified multiple differentially expressed genes (DEGs) (>2-fold change in either direction, p <0.05) (Figure 3), a subset of which were further validated (see below). In addition, Gene Set Enrichment Analysis (GSEA) of DEGs nominated cellular pathways that were significantly enriched in the B^homo^KA compared to KA mice, such as enrichment of fatty acid metabolism and adipogenesis signatures, suggesting altered lipid metabolism as a potential driver of accelerated tumorigenesis observed in the setting of bi-allelic *BAP1* deletions (Figure 3).

**Figure 3:**
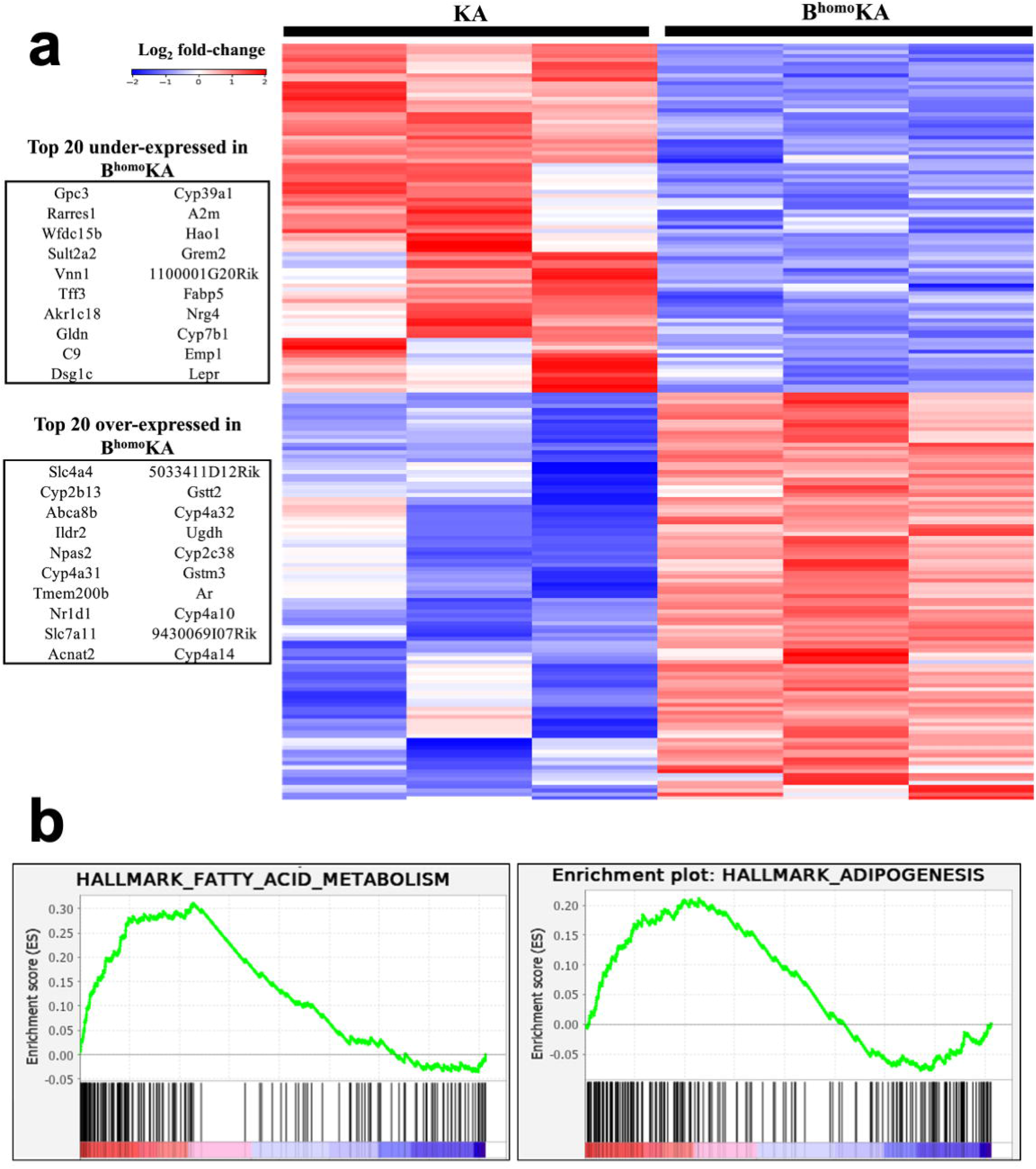
Hepatic tumors from KA and B^homo^KA experimental GEMM cohorts have unique gene expression patterns. (**a**) KA and B^homo^KA liver lesions have differential gene expression by Affymetrix microarray. Supervised clustering heatmap of Affymetrix microarray data (3 KA samples versus 3 B^homo^KA samples, log_2_ fold change >2, p <0.05). The top 20 under- and overexpressed genes in the B^homo^KA cohort are pictured to the left of the heatmap; (**b**) Gene set enrichment analysis (GSEA) of differentially expressed transcripts in microarray of KA and B^homo^KA hepatic tumors shows positive enrichment in hallmark gene signatures for fatty acid metabolism and adipogenesis.

*SLC7A11* was found to be one of the highest over-expressed genes in B^homo^KA tumors compared to KA tumors (Figure 3). Recent studies have highlighted the potential role of altered cystine transport as the mechanism by which loss of BAP1 promotes tumorigenesis, specifically via upregulation of the cystine/glutamate antiporter xCT that is encoded by *SLC7A11* [33]. Therefore, IHC for SLC7A11/xCT expression was performed on harvested liver sections to validate the microarray findings. While KA tumors had minimal to absent SLC7A11/xCT expression within neoplastic tissues, robust SLC7A11/xCT expression was observed in all of the examined B^homo^KA tumors, but particularly so within the HCC foci (Figure 4). Surprisingly, in addition to faint membranous staining, we noted strong granular SLC7A11/xCT staining in the cytoplasm of neoplastic cells of B^homo^KA mice (Figure 4), the significance of which is unclear. Nonetheless, the IHC data confirmed the microarray findings that SLC7A11/xCT was upregulated within B^homo^KA compared to KA tumors, validating prior data that loss of BAP1 upregulates this cystine transporter and regulator of ferroptosis in neoplastic cells [33–34].

**Figure 4:**
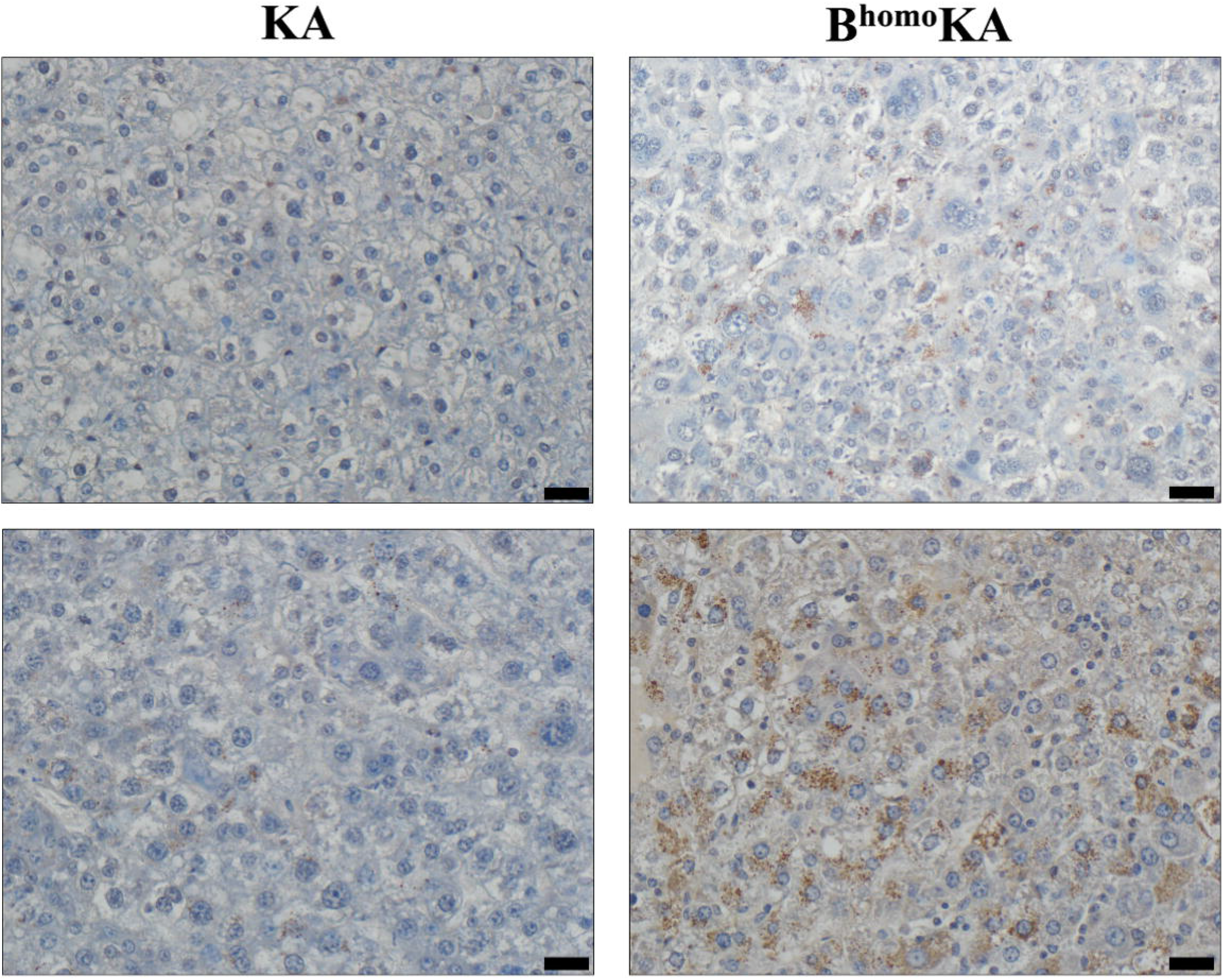
Loss of BAP1 is associated with increased hepatic expression of the cystine/glutamate transporter (SLC7A11/xCT) in GEMM experimental cohorts. Immunohistochemical (IHC) staining for SLC7A11/xCT shows strong staining throughout the neoplastic tissue of B^homo^KA mice as compared to KA, which do not demonstrate staining. Images in the upper panel were taken at 200x magnification. Images in the lower panel were taken at 400x magnification, which more clearly demonstrates the strong granular SLC7A11/xCT cytoplasmic staining in neoplastic cells of B^homo^KA liver sections. Scale bar is 50 μm for 200x images and 25 μm for 400x images.

## Discussion

In this study, we have established a novel autochthonous mouse model of ICC that incorporates bi-allelic *Bap1* deletion in conjunction with expression of mutant *Kras* within the *Albumin*-expressing progenitor population in the developing liver. Of note, our data establishes that bi-allelic *Bap1* deletion is a requirement to elicit an ICC phenotype in the presence of mutant *Kras*, as observed in B^homo^KA animals. Mice that retain even partial Bap1 function (B^het^KA mice) demonstrate pure HCC, without ICC foci, reiterating the importance of this complete loss of function requirement. There have been previous reports of ICC GEMMs, most of which, like our B^homo^KA model, develop combined HCC and ICC. These include models with liver-specific *Pten* and *Smad4* deletion [45], *Kras* activation and *TP53* deletion [35], *Kras* activation and *Pten* deletion [59], *Sav1* deletion [60], and expression of the Notch receptor intracellular domain (*NICD*) [61]. For example, O’Dell and colleagues subsequently demonstrated that mutant *Kras* expression alone led to ICC development with low penetrance and after a long latency, while the combination of mutant *Kras* expression and *Trp53* deletion resulted in near-complete penetrance of ICC development, and often with evidence of metastatic disease, in addition to HCC development [35]. More recently, a mouse model incorporating liver-specific mutant *Kras* expression and *Pten* deletion demonstrated mixed ICC and HCC when heterozygous *Pten* deletion was present, but pure ICC development with *Kras* activation and homozygous *Pten* deletion [59].

Recent molecular studies of human tumor samples have demonstrated that somatic mutations in *BAP1* and *KRAS* are among the most frequent mutations present in ICC [23]. To the best of our knowledge, our model represents the first murine model of ICC to incorporate a *Bap1* deletion. The tumors induced in the B^homo^KA model recapitulate many of the key histologic features of human ICC, including demonstration of the multistep progression of histopathological changes from bile duct hyperplasia to dysplasia, carcinoma *in situ*, and culminating in ICC. Both well-differentiated ICC, characterized by intact glandular architecture and mucin production, and poorly-differentiated ICC with minimal gland formation, large pleomorphic cells, nuclear atypia, and frequent mitoses were seen.

In our model, liver-specific deletion of *Bap1* and *Kras* activation were regulated by Cre recombinase expression driven by the *Albumin* promoter, resulting in Cre-mediated recombination in liver progenitor cells in late embryonic development, as well as in mature hepatocytes and cholangiocytes. Based on morphological and immunohistochemical studies, HCC and ICC were considered to originate from mature hepatic parenchymal cells, hepatocytes and cholangiocytes, respectively [62]. Over time, evidence has accumulated that suggests multiple potential cells of origin of both HCC and ICC. These range from more mature hepatocytes and cholangiocytes, to less-differentiated oval cells and hepatoblasts, and, finally, to the least-differentiated liver stem cells [63–65]. As our model targets liver-specific *Bap1* deletion and oncogenic *Kras* expression to multiple Alb-expressing cell types (an inherent pitfall of the *Alb*-Cre model), the precise cell or cells of origin in our GEMMs remains speculative in the absence of lineage tracing studies. Nonetheless, our finding that ICC only developed in mice with homozygous deletion of *Bap1* and *Kras* activation, suggests that complete loss of *Bap1* cooperates with oncogenic *Kras* to either preferentially drive the putative cell(s) of origin down the pathway of cholangiocyte differentiation and/or transdifferentiation of hepatocytes into biliary-like cells.

Ferroptosis is a metabolic, stress-induced, non-apoptotic form of regulated cell death [33]. The role of ferroptosis in oncogenesis is a topic of ongoing investigation. Recently, BAP1 was found to link ferroptosis to tumor suppression [33]. Specifically, it was found that *Bap1* functions as a tumor suppressor gene by inhibiting cystine uptake into cells, thereby rendering them more sensitive to ferroptosis. It does so by repressing the expression of xCT, via deubiquitination of H2Aub on the *SLC7A11* promoter, which encodes for this protein. xCT is the catalytic subunit of the cystine/glutamate antiporter, the major transporter of extracellular cystine. Our microarray analysis revealed that *SLC7A11* was overexpressed in B^homo^KA tumors as compared to KA tumors. Furthermore, IHC validated increased levels of the xCT protein within the neoplastic lesions found in B^homo^KA livers, compared to minimal expression in KA mice. These results suggest that in B^homo^KA mice, the lack of *SLC7A11* repression by BAP1 might enable malignant cholangiocytes to evade ferroptosis and thereby undergo malignant transformation.

## Conclusions

We describe a novel GEMM for ICC that is driven by liver-specific bi-allelic *Bap1* deletion and expression of oncogenic *Kras*. Our autochthonous model demonstrates that complete loss of Bap1 function is a requirement for developing an ICC phenotype in the setting of mutant *Kras* expression. Continued investigations utilizing our mouse model should provide a better understanding of the pathogenesis of ICC (e.g., markers associated with early changes during ICC initiation, mechanisms underlying tumor progression) and facilitate the development of new, targeted therapies and/or chemoprevention for an increasingly prevalent and persistently fatal disease.

## Supplementary Materials

The following are available online at www.mdpi.com/xxx/s1, Figure S1: Mouse genotyping protocol, Figure S2: Hepatic histopathologic time-lapse of GEMM experimental cohorts, Figure S3: Validation of BAP1 protein expression loss, Figure S4: GEMM experimental cohorts with constitutional KRAS activation develop lung lesions.

## Author Contributions

Conceptualization, A.M., S.G., B.G, and R.M.; methodology, A.M, S.G, B.G., S.F.B., and R.M.; data generation, R.M., S.F.B., A.H., W.C.F., J.Z., J.R.B.; writing— original draft preparation, R.M and S.F.B. All authors have read and agreed to the published version of the manuscript.

## Funding

This research was funded by National Institutes of Health, grant number T32 CA 009599.

## Institutional Review Board Statement

The study was conducted according to the guidelines of National Institutes of Health, and approved by the Institutional Animal Care and Use Committee (IUCUC) of The University of Texas MD Anderson Cancer Center (protocol code 00001937-RN00 l, expiring May 2022).

## Data Availability Statement

The data presented in this study are deposited in NCBI’s Gene Expression Omnibus under GEO Series accession number GSE183554 (https://www.ncbi.nlm.nih.gov/geo/query/acc.cgi?acc=GSE183554). To review GEO accession GSE183554: Go to https://urldefense.com/v3/_https://www.ncbi.nlm.nih.gov/geo/query/acc.cgi?acc=GSE183554__;!!PfbeBCCAmug!yxRfs7maVWvBKsufS2dLRYJ49lon75ndjapD679Ra3fpYnvFB_QvHvgKmgX6AkOQL3vNCQ$. Enter token snelucggljgjtcr into the box.

## Acknowledgments

The authors would like to thank the Advanced Technology Genomics Core (grant CA 16672), Carol Johnson and the MD Anderson Surgery Histology Laboratory, Joshua Dinn, Tamara Griffiths, (Mary) Rose Reisenauer, and Gita Dangol for their assistance with this manuscript.

## Conflicts of Interest

A.M. receives royalties for a pancreatic cancer biomarker test from Cosmos Wisdom Biotechnology, and on a patent that has been licensed by Johns Hopkins University to ThriveEarlier Detection. A.M. also serves as a consultant for Freenome. The remaining authors declare no conflict of interest.

## Notes

https://urldefense.com/v3/__https://www.ncbi.nlm.nih.gov/geo/query/acc.cgi?acc=GSE183554__;!!PfbeBCCAmug!yxRfs7maVWvBKsufS2dLRYJ49lon75ndjapD679Ra3fpYnvFB_QvHvgKmgX6AkOQL3vNCQ$

### Competing Interest Statement

A.M. receives royalties for a pancreatic cancer biomarker test from Cosmos Wisdom Biotechnology, and on a patent that has been licensed by Johns Hopkins University to ThriveEarlier Detection. A.M. also serves as a consultant for Freenome. The remaining authors declare no conflict of interest.

## References

1. Banales JM, Marin JJG, Lamarca A, Rodrigues PM, Khan SA, Roberts LR, et al. Cholangiocarcinoma 2020: the next horizon in mechanisms and management. Nat Rev Gastroenterol Hepatol. 2020 Sep;17(9):557–88.

2. Rizvi S, Gores GJ. Pathogenesis, Diagnosis, and Management of Cholangiocarcinoma. Gastroenterology. 2013 Dec;145(6):1215–29.

3. Dodson RM, Weiss MJ, Cosgrove D, Herman JM, Kamel I, Anders R, et al. Intrahepatic cholangiocarcinoma: management options and emerging therapies. J Am Coll Surg. 2013 Oct;217(4):736–750.e4.

4. Shaib Y, El-Serag HB. The epidemiology of cholangiocarcinoma. Semin Liver Dis. 2004 May;24(2):115–25.

5. Kirstein MM, Vogel A. Epidemiology and Risk Factors of Cholangiocarcinoma. Visc Med. 2016 Dec;32(6):395–400.

6. Siegel RL, Miller KD, Jemal A. Cancer statistics, 2020. CA Cancer J Clin. 2020;70(1):7–30.

7. Shaib YH, Davila JA, McGlynn K, El-Serag HB. Rising incidence of intrahepatic cholangiocarcinoma in the United States: a true increase? J Hepatol. 2004 Mar;40(3):472–7.

8. Saha SK, Zhu AX, Fuchs CS, Brooks GA. Forty Year Trends in Cholangiocarcinoma Incidence in the U.S.: Intrahepatic Disease on the Rise. The Oncologist. 2016 May;21(5):594–9.

9. Rizzo A, Brandi G. BILCAP trial and adjuvant capecitabine in resectable biliary tract cancer: reflections on a standard of care. Expert Rev Gastroenterol Hepatol. 2020 Dec 18;1–3.

10. Utuama O, Permuth JB, Dagne G, Sanchez-Anguiano A, Alman A, Kumar A, et al. Neoadjuvant Chemotherapy for Intrahepatic Cholangiocarcinoma: A Propensity Score Survival Analysis Supporting Use in Patients with High-Risk Disease. Ann Surg Oncol. 2021 Jan 7;

11. Buettner S, van Vugt JL, IJzermans JN, Groot Koerkamp B. Intrahepatic cholangiocarcinoma: current perspectives. OncoTargets Ther. 2017;10:1131–42.

12. Lamarca A, Ross P, Wasan HS, Hubner RA, McNamara MG, Lopes A, et al. Advanced Intrahepatic Cholangiocarcinoma: Post Hoc Analysis of the ABC-01, -02, and -03 Clinical Trials. J Natl Cancer Inst. 2020 Feb 1;112(2):200–10.

13. Kish M, Chan K, Perry K, Ko YJ. A systematic review and network meta-analysis of adjuvant therapy for curatively resected biliary tract cancers. Curr Oncol Tor Ont. 2020 Feb;27(1):e20–6.

14. Lamarca A, Edeline J, McNamara MG, Hubner RA, Nagino M, Bridgewater J, et al. Current standards and future perspectives in adjuvant treatment for biliary tract cancers. Cancer Treat Rev. 2020 Mar;84:101936.

15. Ebata T, Hirano S, Konishi M, Uesaka K, Tsuchiya Y, Ohtsuka M, et al. Randomized clinical trial of adjuvant gemcitabine chemotherapy versus observation in resected bile duct cancer. Br J Surg. 2018 Feb;105(3):192–202.

16. Edeline J, Benabdelghani M, Bertaut A, Watelet J, Hammel P, Joly J-P, et al. Gemcitabine and Oxaliplatin Chemotherapy or Surveillance in Resected Biliary Tract Cancer (PRODIGE 12-ACCORD 18-UNICANCER GI): A Randomized Phase III Study. J Clin Oncol Off J Am Soc Clin Oncol. 2019 Mar 10;37(8):658–67.

17. Malka D, Edeline J. Adjuvant capecitabine in biliary tract cancer: a standard option? Lancet Oncol. 2019 May;20(5):606–8.

18. Primrose JN, Fox RP, Palmer DH, Malik HZ, Prasad R, Mirza D, et al. Capecitabine compared with observation in resected biliary tract cancer (BILCAP): a randomised, controlled, multicentre, phase 3 study. Lancet Oncol. 2019 May;20(5):663–73.

19. Shroff RT, Kennedy EB, Bachini M, Bekaii-Saab T, Crane C, Edeline J, et al. Adjuvant Therapy for Resected Biliary Tract Cancer: ASCO Clinical Practice Guideline. J Clin Oncol Off J Am Soc Clin Oncol. 2019 Apr 20;37(12):1015–27.

20. Banales JM, Cardinale V, Carpino G, Marzioni M, Andersen JB, Invernizzi P, et al. Expert consensus document: Cholangiocarcinoma: current knowledge and future perspectives consensus statement from the European Network for the Study of Cholangiocarcinoma (ENS-CCA). Nat Rev Gastroenterol Hepatol. 2016 May;13(5):261–80.

21. Hu L-S, Zhang X-F, Weiss M, Popescu I, Marques HP, Aldrighetti L, et al. Recurrence Patterns and Timing Courses Following Curative-Intent Resection for Intrahepatic Cholangiocarcinoma. Ann Surg Oncol. 2019 Aug;26(8):2549–57.

22. Bagante F, Spolverato G, Weiss M, Alexandrescu S, Marques HP, Aldrighetti L, et al. Defining Long-Term Survivors Following Resection of Intrahepatic Cholangiocarcinoma. J Gastrointest Surg Off J Soc Surg Aliment Tract. 2017 Nov;21(11):1888–97.

23. Farshidfar F, Zheng S, Gingras M-C, Newton Y, Shih J, Robertson AG, et al. Integrative Genomic Analysis of Cholangiocarcinoma Identifies Distinct IDH-Mutant Molecular Profiles. Cell Rep. 2017 Mar;18(11):2780–94.

24. Chan-on W, Nairismägi M-L, Ong CK, Lim WK, Dima S, Pairojkul C, et al. Exome sequencing identifies distinct mutational patterns in liver fluke–related and non-infection-related bile duct cancers. Nat Genet. 2013 Dec;45(12):1474–8.

25. Johnson L, Mercer K, Greenbaum D, Bronson RT, Crowley D, Tuveson DA, et al. Somatic activation of the K-ras oncogene causes early onset lung cancer in mice. Nature. 2001 Apr;410(6832):1111–6.

26. Li S, Balmain A, Counter CM. A model for RAS mutation patterns in cancers: finding the sweet spot. Nat Rev Cancer. 2018 Dec;18(12):767–77.

27. Carbone M, Yang H, Pass HI, Krausz T, Testa JR, Gaudino G. BAP1 and cancer. Nat Rev Cancer. 2013 Mar;13(3):153–9.

28. Ventii KH, Devi NS, Friedrich KL, Chernova TA, Tighiouart M, Van Meir EG, et al. BRCA1-associated protein-1 is a tumor suppressor that requires deubiquitinating activity and nuclear localization. Cancer Res. 2008 Sep 1;68(17):6953–62.

29. Luchini C, Veronese N, Yachida S, Cheng L, Nottegar A, Stubbs B, et al. Different prognostic roles of tumor suppressor gene BAP1 in cancer: A systematic review with meta-analysis. Genes Chromosomes Cancer. 2016 Oct;55(10):741–9.

30. Peña-Llopis S, Vega-Rubín-de-Celis S, Liao A, Leng N, Pavía-Jiménez A, Wang S, et al. BAP1 loss defines a new class of renal cell carcinoma. Nat Genet. 2012 Jun 10;44(7):751–9.

31. Qin J, Zhou Z, Chen W, Wang C, Zhang H, Ge G, et al. BAP1 promotes breast cancer cell proliferation and metastasis by deubiquitinating KLF5. Nat Commun. 2015 Dec;6(1):8471.

32. Testa JR, Cheung M, Pei J, Below JE, Tan Y, Sementino E, et al. Germline BAP1 mutations predispose to malignant mesothelioma. Nat Genet. 2011 Aug 28;43(10):1022–5.

33. Zhang Y, Shi J, Liu X, Feng L, Gong Z, Koppula P, et al. BAP1 links metabolic regulation of ferroptosis to tumor suppression. Nat Cell Biol. 2018 Oct;20(10):1181–92.

34. Zhang Y, Koppula P, Gan B. Regulation of H2A ubiquitination and SLC7A11 expression by BAP1 and PRC1. Cell Cycle Georget Tex. 2019 Apr;18(8):773–83.

35. O’Dell MR, Li Huang J, Whitney-Miller CL, Deshpande V, Rothberg P, Grose V, et al. *Kras G12D* and *p53* Mutation Cause Primary Intrahepatic Cholangiocarcinoma. Cancer Res. 2012 Mar 15;72(6):1557–67.

36. Postic C, Magnuson MA. DNA excision in liver by an albumin-Cre transgene occurs progressively with age.

37. Gu Y-F, Cohn S, Christie A, McKenzie T, Wolff N, Do QN, et al. Modeling Renal Cell Carcinoma in Mice: Bap1 and Pbrm1 Inactivation Drive Tumor Grade. Cancer Discov. 2017 Aug;7(8):900–17.

38. Wang S-S, Gu Y-F, Wolff N, Stefanius K, Christie A, Dey A, et al. Bap1 is essential for kidney function and cooperates with Vhl in renal tumorigenesis. Proc Natl Acad Sci U S A. 2014 Nov 18;111(46):16538–43.

39. Kühn R, Schwenk F, Aguet M, Rajewsky K. Inducible Gene Targeting in Mice. Sci New Ser. 1995;269(5229):1427–9.

40. Jackson EL, Willis N, Mercer K, Bronson RT, Crowley D, Montoya R, et al. Analysis of lung tumor initiation and progression using conditional expression of oncogenic K-ras. Genes Dev. 2001 Dec 15;15(24):3243–8.

41. Gupta S, Pramanik D, Mukherjee R, Campbell NR, Elumalai S, de Wilde RF, et al. Molecular determinants of retinoic acid sensitivity in pancreatic cancer. Clin Cancer Res Off J Am Assoc Cancer Res. 2012 Jan 1;18(1):280–9.

42. Carvalho BS, Irizarry RA. A framework for oligonucleotide microarray preprocessing. Bioinforma Oxf Engl. 2010 Oct 1;26(19):2363–7.

43. Ritchie ME, Phipson B, Wu D, Hu Y, Law CW, Shi W, et al. limma powers differential expression analyses for RNA-sequencing and microarray studies. Nucleic Acids Res. 2015 Apr 20;43(7):e47.

44. Subramanian A, Tamayo P, Mootha VK, Mukherjee S, Ebert BL, Gillette MA, et al. Gene set enrichment analysis: a knowledge-based approach for interpreting genome-wide expression profiles. Proc Natl Acad Sci U S A. 2005 Oct 25;102(43):15545–50.

45. Xu X. Induction of intrahepatic cholangiocellular carcinoma by liver-specific disruption ofSmad4 andPten in mice. J Clin Invest. 2006 Jul 3;116(7):1843–52.

46. Kenerson HL, Yeh MM, Kazami M, Jiang X, Riehle KJ, McIntyre RL, et al. Akt and mTORC1 have different roles during liver tumorigenesis in mice. Gastroenterology. 2013 May;144(5):1055–65.

47. Saborowski A, Saborowski M, Davare MA, Druker BJ, Klimstra DS, Lowe SW. Mouse model of intrahepatic cholangiocarcinoma validates FIG-ROS as a potent fusion oncogene and therapeutic target. Proc Natl Acad Sci U S A. 2013 Nov 26;110(48):19513–8.

48. Dey A, Seshasayee D, Noubade R, French DM, Liu J, Chaurushiya MS, et al. Loss of the tumor suppressor BAP1 causes myeloid transformation. Science. 2012 Sep 21;337(6101):1541–6.

49. Balaton AJ, Nehama-Sibony M, Gotheil C, Callard P, Baviera EE. Distinction between hepatocellular carcinoma, cholangiocarcinoma, and metastatic carcinoma based on immunohistochemical staining for carcinoembryonic antigen and for cytokeratin 19 on paraffin sections. J Pathol. 1988 Dec;156(4):305–10.

50. Johnson DE, Herndier BG, Medeiros LJ, Warnke RA, Rouse RV. The diagnostic utility of the keratin profiles of hepatocellular carcinoma and cholangiocarcinoma. Am J Surg Pathol. 1988 Mar;12(3):187–97.

51. Chu PG, Ishizawa S, Wu E, Weiss LM. Hepatocyte antigen as a marker of hepatocellular carcinoma: an immunohistochemical comparison to carcinoembryonic antigen, CD10, and alphafetoprotein. Am J Surg Pathol. 2002 Aug;26(8):978–88.

52. Lau SK, Prakash S, Geller SA, Alsabeh R. Comparative immunohistochemical profile of hepatocellular carcinoma, cholangiocarcinoma, and metastatic adenocarcinoma. Hum Pathol. 2002 Dec;33(12):1175–81.

53. Fan Z, van de Rijn M, Montgomery K, Rouse RV. Hep par 1 antibody stain for the differential diagnosis of hepatocellular carcinoma: 676 tumors tested using tissue microarrays and conventional tissue sections. Mod Pathol Off J U S Can Acad Pathol Inc. 2003 Feb;16(2):137–44.

54. Ryu HS, Lee K, Shin E, Kim SH, Jing J, Jung HY, et al. Comparative Analysis of Immunohistochemical Markers for Differential Diagnosis of Hepatocelluar Carcinoma and Cholangiocarcinoma. Tumori J. 2012 Jul;98(4):478–84.

55. Gagliano EF. Reticulin stain in the fine needle aspiration differential diagnosis of liver nodules. Acta Cytol. 1995 Jun;39(3):596–8.

56. Bergman S, Graeme-Cook F, Pitman MB. The usefulness of the reticulin stain in the differential diagnosis of liver nodules on fine-needle aspiration biopsy cell block preparations. Mod Pathol Off J U S Can Acad Pathol Inc. 1997 Dec;10(12):1258–64.

57. Wee A. Fine needle aspiration biopsy of the liver: Algorithmic approach and current issues in the diagnosis of hepatocellular carcinoma. CytoJournal. 2005 Jun 8;2:7.

58. Nahon JL, Tratner I, Poliard A, Presse F, Poiret M, Gal A, et al. Albumin and alpha-fetoprotein gene expression in various nonhepatic rat tissues. J Biol Chem. 1988 Aug 15;263(23):11436–42.

59. Ikenoue T, Terakado Y, Nakagawa H, Hikiba Y, Fujii T, Matsubara D, et al. A novel mouse model of intrahepatic cholangiocarcinoma induced by liver-specific Kras activation and Pten deletion. Sci Rep. 2016 Apr 29;6(1):23899.

60. Lee K-P, Lee J-H, Kim T-S, Kim T-H, Park H-D, Byun J-S, et al. The Hippo–Salvador pathway restrains hepatic oval cell proliferation, liver size, and liver tumorigenesis. Proc Natl Acad Sci U S A. 2010 May 4;107(18):8248–53.

61. Zender S, Nickeleit I, Wuestefeld T, Sörensen I, Dauch D, Bozko P, et al. A critical role for notch signaling in the formation of cholangiocellular carcinomas. Cancer Cell. 2013 Jun 10;23(6):784–95.

62. Yamamoto M, Xin B, Watanabe K, Ooshio T, Fujii K, Chen X, et al. Oncogenic Determination of a Broad Spectrum of Phenotypes of Hepatocyte-Derived Mouse Liver Tumors. Am J Pathol. 2017 Dec;187(12):2711–25.

63. Fausto N. Liver regeneration and repair: hepatocytes, progenitor cells, and stem cells. Hepatol Baltim Md. 2004 Jun;39(6):1477–87.

64. Oikawa T. Cancer Stem cells and their cellular origins in primary liver and biliary tract cancers. Hepatology. 2016 Aug;64(2):645–51.

65. Sia D, Villanueva A, Friedman SL, Llovet JM. Liver Cancer Cell of Origin, Molecular Class, and Effects on Patient Prognosis. Gastroenterology. 2017 Mar;152(4):745–61.

