## Supplementary Figure Legends for "Oncogenic KRAS Requires Complete Loss of BAP1 Function for Development of Murine Intrahepatic Cholangiocarcinoma"

**Supplementary Figure 1: Mouse genotyping protocol.** Primer sequences for mouse genes examined (**a**) and representative agarose gel electrophoresis with expected amplicon sizes (**b**).

**Supplementary Figure 2: Hepatic histopathologic time-lapse of GEMM experimental cohorts.** Disease progression in KA (**a**), $B^{homo}A$ (**b**), and $B^{het}KA$ (**c**), and $B^{homo}KA$ mice (**d**). Images were taken with 200x magnification. Scale bar is 50 μm.

**Supplementary Figure 3: Validation of BAP1 protein expression loss.** BAP1 staining of experimental cohorts shows loss of expression in hepatic parenchyma (**a**) and primary liver tumors (**b**). Images were taken with 200x magnification. Scale bar is 50 μ*m*.

**Supplementary Figure 4: GEMM experimental cohorts with constitutional Kras activation develop lung lesions.** KA and $B^{het}KA$ mice exhibit well-circumscribed lung lesions (**a**) that stain diffusely for CK-19 but not for Hep Par 1, most consistent with primary lung adenocarcinoma (**b**). H&E images were taken with 100x magnification and IHC with 200x. Scale bar is 100 μ*m* for H&E images and 50 μ*m* for IHC.
