## Supplementary Figures 1-4 for "Oncogenic KRAS Requires Complete Loss of BAP1 Function for Development of Murine Intrahepatic Cholangiocarcinoma"

**a**

| Gene | Primer Sequences |
|---|---|
| Tg(Alb-cre) | 5’TGCAAACATCACATGCACAC3’ |
| | 5’GAAGCAGAAGCTTAGGAAGATG3’ |
| | 5’TTGGCCCCTTACCATAACTG3’ |
| BAP1 | 5'GGGCACATCTGATCCTCAGAGCTA3' |
| | 5'GGCAGTGGTGGCAAATGAGACCTT3' |
| | 5'GCACTGACAGCTGCCCATCTGAA3' |
| KRAS | 5'GTCTTTCCCCAGCACAGTGC3' |
| | 5'CTCTTGCCTACGCCACCAGCTC3' |
| | 5AGCTAGCCACCATGGCTTGAGTAAGTCTGCA3’ |

**b**

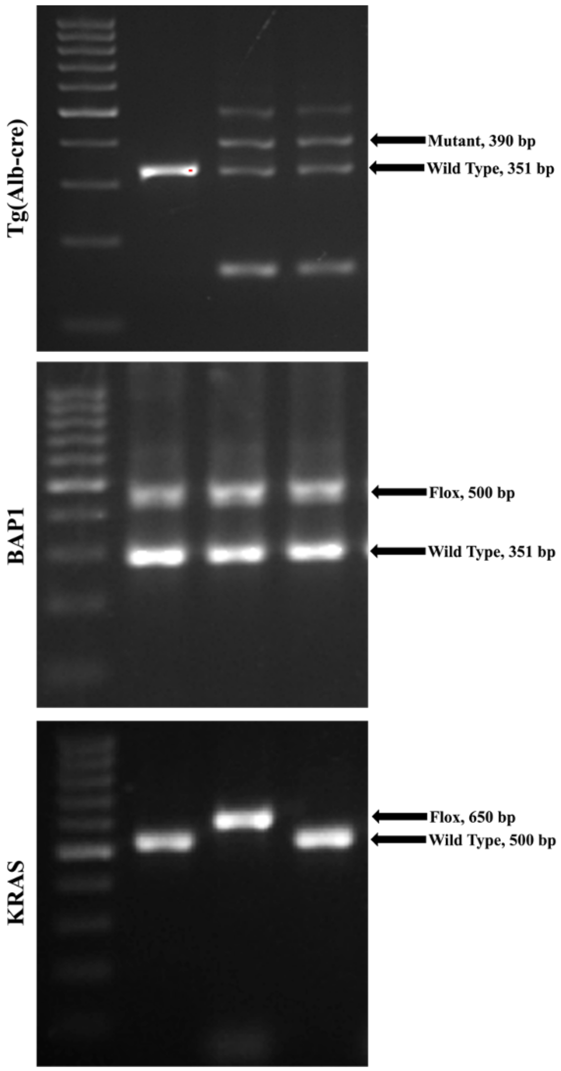


**Supplementary Figure 1: Mouse genotyping protocol.** Primer sequences for mouse genes examined (**a**) and representative agarose gel electrophoresis with expected amplicon sizes (**b**).

**a**

**KA**

| Age (weeks) | 4 - 8 | 12 - 16 | 20 - 28 | 32 - 40 | 44 - 60 |
|---|---|---|---|---|---|
| Histology | • Normal liver | • Fatty metamorphosis of hepatocytes<br>• Microvesicular steatosis | • Fatty metamorphosis of hepatocytes<br>• Micro- and macrovesicular steatosis<br>• Liver parenchyma congestion<br>• Hepatic adenomas | • Fatty metamorphosis of hepatocytes<br>• Micro- and macrovesicular steatosis<br>• Liver parenchyma congestion<br>• Hepatic adenomas<br>• Well-differentiated to advanced HCC | • Fatty metamorphosis of hepatocytes<br>• Micro- and macrovesicular steatosis<br>• Liver parenchyma congestion<br>• Hepatic adenomas<br>• Well-differentiated to advanced HCC |

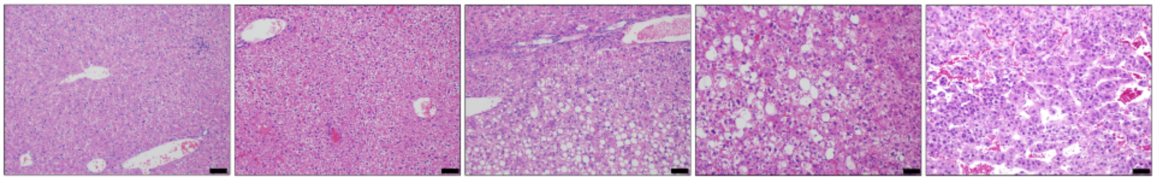

**b**

**$B^{homo}A$**

| Age (weeks) | 4 | 8 - 20 | 24 - 32 | 36 - 40 | 44 - 52 |
|---|---|---|---|---|---|
| Histology | • Normal liver | • Enlarged hepatocytes | • Enlarged hepatocytes<br>• Fatty metamorphosis of hepatocytes<br>• Liver parenchyma congestion<br>• Rare hepatic adenomas | • Enlarged hepatocytes<br>• Fatty metamorphosis of hepatocytes<br>• Liver parenchyma congestion<br>• Rare hepatic adenomas<br>• Rare well-differentiated HCC | • Enlarged hepatocytes<br>• Fatty metamorphosis of hepatocytes<br>• Liver parenchyma congestion<br>• Hepatic adenomas<br>• Well-differentiated and rare advanced HCC |

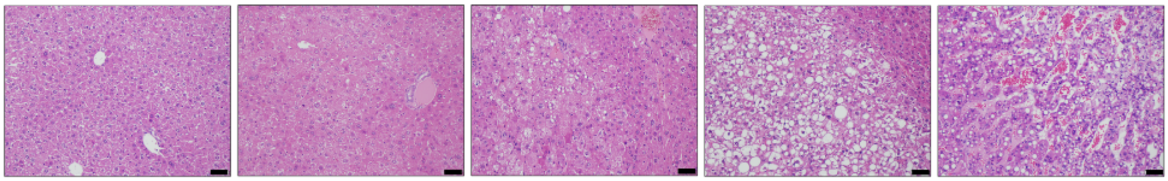

**c**

**$B^{het}KA$**

| Age (weeks) | 4 - 16 | 20 - 24 | 28 - 32 | 36 - 40 | 44 - 56 |
|---|---|---|---|---|---|
| Histology | • Normal hepatocytes<br>• Dilated blood vessels | • Normal hepatocytes<br>• Dilated blood vessels<br>• Hepatic adenomas | • Dilated blood vessels<br>• Hepatic adenomas<br>• Well-differentiated HCC | • Dilated blood vessels<br>• Hepatic adenomas<br>• Well-differentiated and advanced HCC | • Dilated blood vessels<br>• Hepatic adenomas<br>• Well-differentiated and advanced HCC |

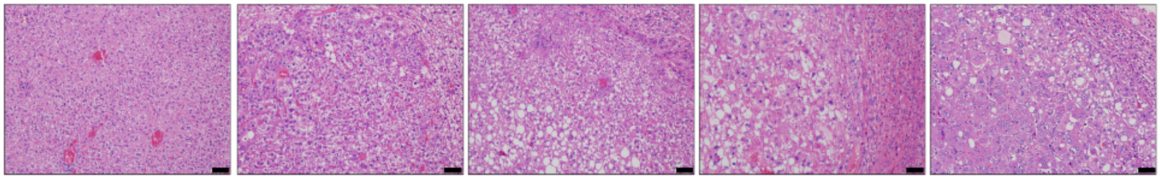

**d**

**$B^{homo}KA$**

| Age (weeks) | 4 | 8 | 12 - 16 | 20 | 24 |
|---|---|---|---|---|---|
| Histology | • Fatty metamorphosis of hepatocytes<br>• Microvesicular steatosis<br>• Steatohepatis<br>• Liver parenchyma congestion | • Fatty metamorphosis of hepatocytes<br>• Microvesicular steatosis<br>• Steatohepatis<br>• Liver parenchyma congestion<br>• Biliary hyperplasia | • Fatty metamorphosis of hepatocytes<br>• Microvesicular steatosis<br>• Steatohepatis<br>• Liver parenchyma congestion<br>• Biliary hyperplasia<br>• Hepatic adenomas<br>• Well-differentiated HCC | • Fatty metamorphosis of hepatocytes<br>• Microvesicular steatosis<br>• Steatohepatis<br>• Liver parenchyma congestion<br>• Biliary hyperplasia<br>• Hepatic adenomas<br>• Well-differentiated and advanced HCC<br>• Well-differentiated ICC | • Fatty metamorphosis of hepatocytes<br>• Microvesicular steatosis<br>• Steatohepatis<br>• Liver parenchyma congestion<br>• Biliary hyperplasia<br>• Hepatic adenomas<br>• Well-differentiated and advanced HCC<br>• Well-differentiated and advanced ICC |

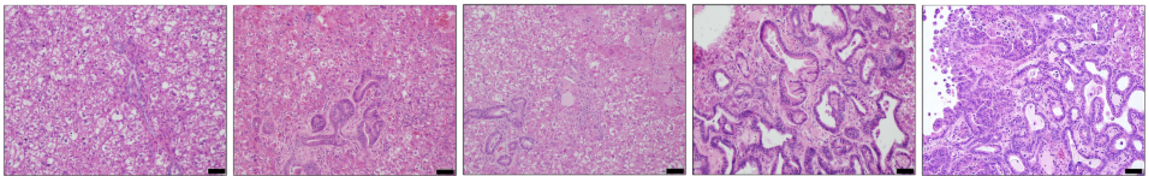

**Supplementary Figure 2: Hepatic histopathologic time-lapse of GEMM experimental cohorts.** Disease progression in KA (**a**), $B^{homo}$A (**b**), and $B^{het}$KA (**c**), and $B^{homo}$KA mice (**d**). Images were taken with 200x magnification. Scale bar is 50 μm.

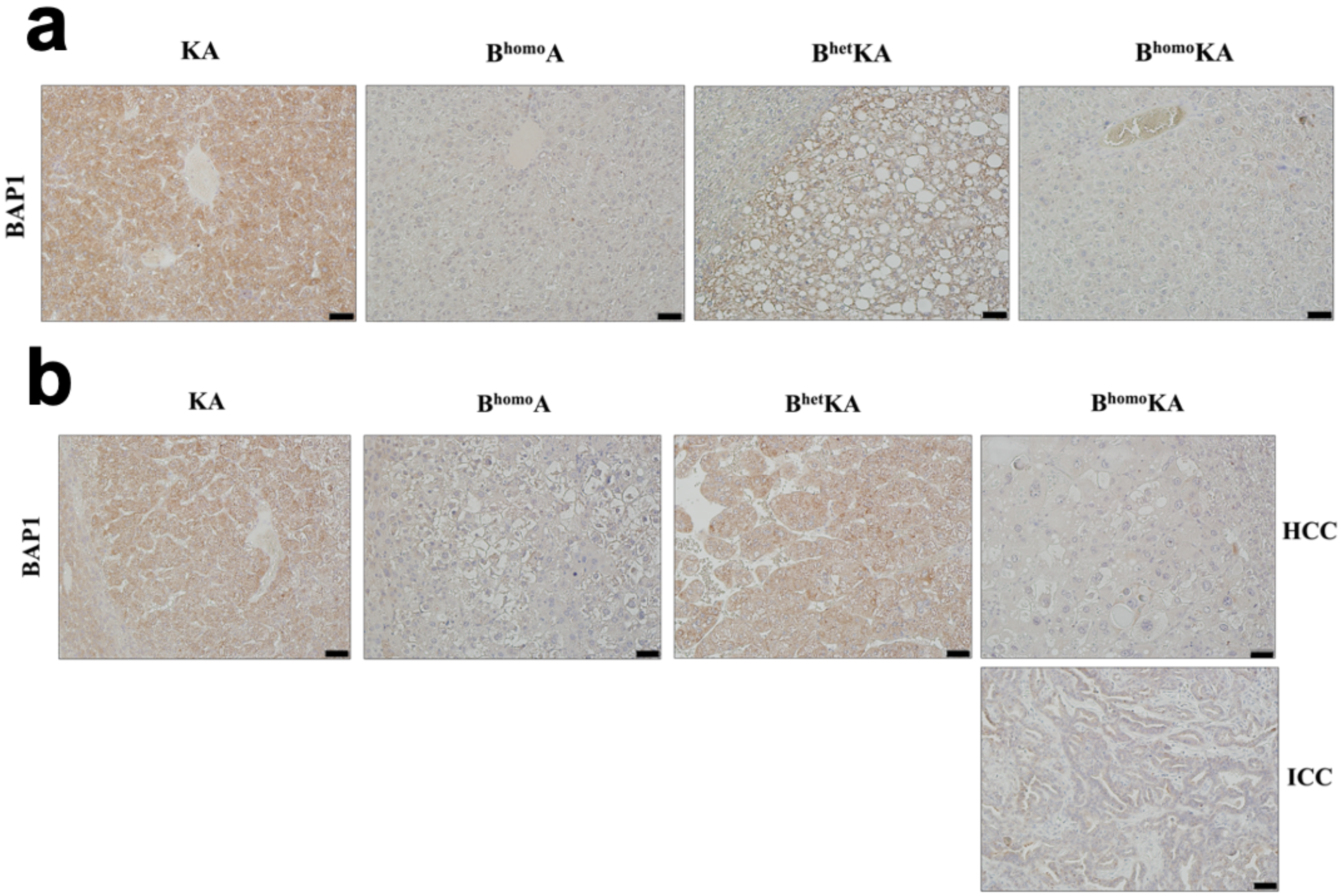


**Supplementary Figure 3: Validation of BAP1 protein expression loss.** BAP1 staining of experimental cohorts shows loss of expression in hepatic parenchyma (**a**) and primary liver tumors (**b**). Images were taken with 200x magnification. Scale bar is 50 μ*m*.

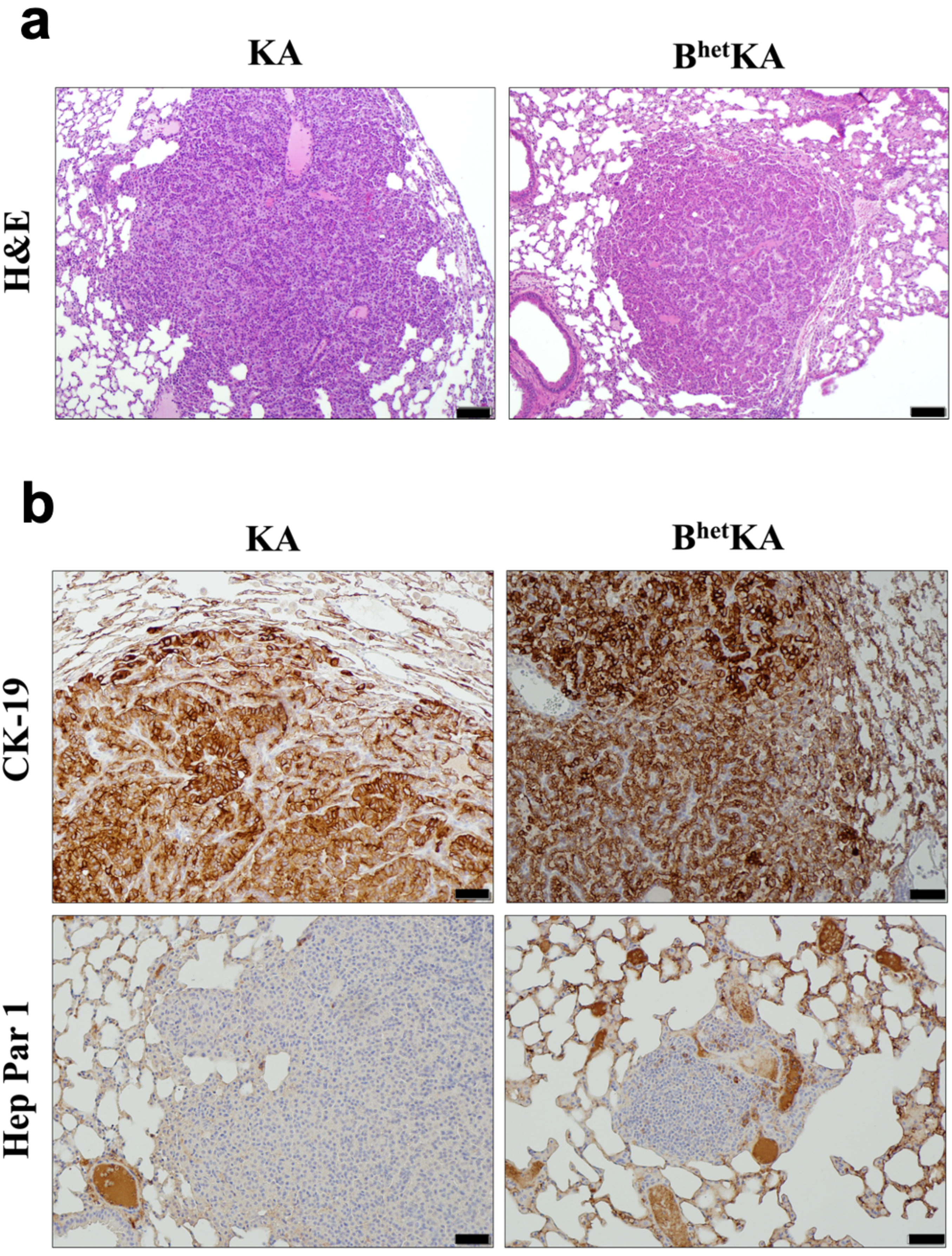
a
KA
B^het KA
H&E
b
KA
B^het KA
CK-19
Hep Par 1

**Supplementary Figure 4: GEMM experimental cohorts with constitutional Kras activation develop lung lesions.** KA and $B^{het}$KA mice exhibit well-circumscribed lung lesions (**a**) that stain diffusely for CK-19 but not for Hep Par 1, most consistent with primary lung adenocarcinoma (**b**). H&E images were taken with 100x magnification and IHC with 200x. Scale bar is 100 $\mu m$ for H&E images and 50 $\mu m$ for IHC.
